# Projecting social contact matrices to different demographic structures

**DOI:** 10.1101/343491

**Authors:** Sergio Arregui, Alberto Aleta, Joaquín Sanz, Yamir Moreno

## Abstract

The modeling of large-scale communicable epidemics has greatly benefited in the last years from the increasing availability of highly detailed data. Particularly, in order to achieve quantitative descriptions of the evolution of epidemics, contact networks and mixing patterns are key. These heterogeneous patterns depend on several factors such as location, socioeconomic conditions, time, and age. This last factor has been shown to encapsulate a large fraction of the observed inter-individual variation in contact patterns, an observation validated by different measurements of age-dependent contact matrices. Recently, several works have studied how to project those matrices to areas where empiric data is not available. However, the dependence of contact matrices on demographic structures and their time evolution has been largely neglected. In this work, we tackle the problem of how to transform an empirical contact matrix that has been obtained for a given demographic structure into a different contact matrix that is compatible with a different demography. The methodology discussed here allows extrapolating a contact structure measured in a particular area to any other whose demographic structure is known, as well as to obtain the time evolution of contact matrices as a function of the demographic dynamics of the populations they refer to. To quantify the effect of considering time-dynamics of contact patterns on disease modeling, we implemented a Susceptible-Exposed-Infected-Recovered (SEIR) model on 16 different countries and evaluated the impact of neglecting the temporal evolution of mixing patterns. Our results show that simulated disease incidence rates, both at the aggregated and age-specific levels, are significantly dependent on contact structures variation driven by demographic evolution. The present work opens the path to eliminate technical biases from model-based impact evaluations of future epidemic threats and warns against the use of contact matrices to model diseases without correcting for demographic evolution or geographic variations.

**Author summary:** Large scale epidemic outbreaks represent an ever increasing threat to humankind. In order to anticipate eventual pandemics, mathematical modeling should not only have the capacity to model in real time an ongoing disease, but also to predict the evolution of potential outbreaks in different locations and times. To this end, computational frameworks need to incorporate, among other ingredients, realistic contact patterns into the models. This not only implies anticipating the demographic structure of the populations under study, but also understanding how demographic evolution reshapes social mixing patterns along time. Here we present a mathematical framework to solve this problem and test our modeling approach on 16 different empirical contact matrices. We also evaluate the impact of an eventual future outbreak by simulating a SEIR scenario in the countries analyzed. Our results show that using outdated or imported contact matrices that do not take into account demographic structure or its evolution can lead to largely misleading conclusions.

## Introduction

During recent years, models on disease transmission have improved in complexity and depth, integrating high-resolution data on demography, mobility and social behavior [1, 2]. Specifically, the topology of social contacts plays a major role in state-of-the-art modeling [3–8]. The complete knowledge of the network of contacts through which an epidemic spreads is usually unreachable or impossible to implement, and for modeling purposes it is useful to remain at the coarse level of age-groups. Under this view, the population under study is divided into different groups according to its age distribution and different contact rates are assumed among these groups. Age-dependent contact patterns give powerful insights on the transmission of diseases where epidemiological risk is correlated to age, either as a result of behavioral or physiological factors. Relevant examples are influenza-like diseases [6–10], pertussis [11], tuberculosis [12, 13], and varicella [14]. Furthermore, they are instrumental for modeling and implementing more efficient interventions [15, 16].

Given the utmost importance of contact heterogeneities, the study of age-dependent social mixing has become a priority in the field. In 2008, Mossong et al. [17] published a seminal work with the measurements of age-dependent contact rates in eight European countries (Belgium, Finland, Germany, Great Britain, Italy, Luxembourg, Netherlands and Poland) via contact diaries. Due to the high cost of gathering empirical data on social contacts, Fumanelli et al. [18] proposed an alternative path consisting on building synthetic contact patterns via the modeling of virtual populations. Nevertheless, other authors have followed the original route open by Mossong et al., measuring empirically the age-dependent social contacts of other countries such as China [19], France [20], Hong-Kong [21], Japan [22], Kenya [23], Russia [24], Uganda [25] and Zimbabwe [26], thus expanding significantly the available data on social mixing in the last few years. In these studies, participants are asked how many contacts they have during a day and with whom. This allows to obtain the (average) number of contacts that an individual of a particular age *i* has with individuals of age-group *j.* The resulting matrix is not symmetric due to the different number of individuals in each age-group. However, it is precisely the demographic structure what imposes constraints in the entries of this matrix, as reciprocity of contacts should be fulfilled at any time (i.e., the total number of contacts reported by age-group *i* with age-group *j* should be ideally equal in the opposite direction). Therefore, an empirical contact matrix, that has been measured on a specific population, should not be used directly, without further considerations, in another population with a different demographic structure.

This issue has important consequences in the field of disease modeling. As contact matrices play a key role in disease forecast, it is essential to assure that the matrices implemented are adapted to the demographic structure of the population considered in order to avoid biased estimations. For some short-cycle diseases like influenza, the time scale of the epidemic is much shorter than the typical times needed for a demographic structure to evolve. That means that, typically, the demographic structure can be safely considered constant [10], and the eventual evolution of the contact matrix can be neglected throughout the simulation of an outbreak. For these diseases, the problems might arise when modelers use contact matrices that are not up to date -for instance, one might wonder whether the patterns reported in [17] in 2008 can be used nowadays, a decade later, during which all the European countries analyzed in that study aged significantly. The same issue appears when a contact matrix measured in a given location (e.g., a specific country) is directly used to simulate disease spreading in another region or country with a different population structure.

The previous considerations are even more troublesome for the case of persistent diseases that need long-term simulations, for which the hypothesis of constant demographic structures does not hold anymore [12]. In those cases, contact matrices should continuously evolve during the simulation to reflect the effect that an evolving demography should exert on contact structures. Furthermore, it remains to be known to what extent the variations between contact matrices coming from different countries are due to differences in the demographic structures, divergent cultural traits and/or methodological differences between studies. For instance, elderly people exhibit higher contact rates with children in African countries than in Europe [26]. This could be explained by the different demographic structures: one might expect to observe higher contact rates toward the younger age strata in Africa than in Europe because their populations have a higher density of young individuals. However, it is not clear yet whether the demographic structure is the only driver of geographical heterogeneity between empirical contact matrices.

The main focus of this work is to study how age contact matrices, originally obtained for a specific setting (country and year), can be adapted to different demographic structures -i.e., to another (location and/or time) setting. To this end, we first study the magnitude of the reciprocity error incurred when the adaptation of empirical social contacts to different age structures is ignored, thus justifying the need of studying possible projections that solve this problem. Next, we analyze different methods to perform these adaptations, highlighting the differences induced in the contact patterns by the use of these methods. We also compare empirical contact matrices of 16 countries in different areas worldwide filtering the influence of the demographic structure. This allows to isolate what are the differences that are caused by other factors such as cultural traits. Finally, we implement a Susceptible-Exposed-Infected-Recovered (SEIR) dynamics to study the differences in prospected incidences that arise when applying the methods analyzed to project social contact matrices.

## Materials and Methods

### Collection of empirical survey matrices

For this work we have gathered 16 different contact matrices coming from several countries: 8 from the POLYMOD project [17] (Belgium, Finland, Germany, Great Britain, Italy, Luxembourg, Netherlands and Poland), China [19], France [20], Hong-Kong [21], Japan [22], Kenya [23], Russia [24], Uganda [25] and Zimbabwe [26].

There are some methodological differences between these studies, thus some pre-processing to homogenize the matrices is required. Specifically, we need to transform them to the same definition of contact matrix and adapt them to the same age-groups. Once this is done, we perform a reciprocity correction (valid for the demographic structure corresponding to the country and year where the survey took place), and we normalize the matrices so that the mean connectivity is equal to one. Details can be found in the Supplementary Information.

### Demographic data

Data regarding the time evolution of demographic structures, either observed in the past or projected until 2050, has been retrieved from the UN population division database [27].

### Projections of a Contact Matrix

The basic problem explored in this work is: how can we transform the (empirical) contact matrix *M_i,j_,* that has been measured for a specific demographic structure *N_i,j_,* into a different contact matrix 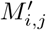 that is compatible with a different demographic structure 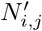? This could mean to adapt data obtained in one specific country to another different region that has a different demography. But the problem can appear even if we remain in the same geographical setting, as a contact matrix measured at a specific time *τ*, could not be valid for an arbitrary time *t* if the demographic structure of that population has changed. In the following sections, we formulate the problem of non-reciprocity and we present and discuss different methods of using contact matrices in an arbitrary demographic structure.

### Method 0 (M0): Unadapted Contact Matrix. The problem of non-reciprocity

We will call *M_i,j_* to the mean number of contacts that an individual of age *i* has with other individuals of age *j* during a certain period of time. This is the magnitude that is usually reported when contact patterns are measured empirically [17, 19–24]. The number of contacts must fulfil reciprocity, i.e., there is the same number of total contacts from age-group *i* to age-group *j* than from *j* to *i.* This imposes the following closure relation for the contact matrix:

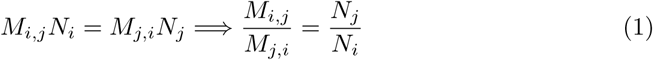

where *N_i_* is the number of individuals of age-group *i.*

Therefore, in the case of an evolving demographic structure for which the ratio 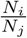 is not constant; the contact matrix *M_i,j_* must change with time. Otherwise we will have non-reciprocal contacts (contacts that inconsistently appear in one direction but not in the other). When comparing different methods for correcting for reciprocity we will usually also compare with the case in which this problem is completely ignored, and the matrix *M_i,j_* is taken directly from the survey without any further consideration. We will refer to this case as Method 0 (M0).

The following methods correct this problem, introducing different transformations of the original contact matrix *M_i,j_,* that was measured in a demographic structure *N_i_,* into a new contact matrix 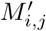 that is well adapted to a new demographic structure 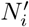 (at least avoiding the problem of no reciprocity).

### Method 1 (M1): Pair-wise correction

The basic problem that we want to avoid is to have a different number of contacts measured from *i* to *j* than from *j* to *i.* Thus, an immediate correction would be to simply average those numbers, so the excess of contacts measured in one direction is transferred to the reciprocal direction. This correction can be formulated as:

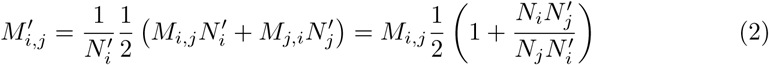

### Method 2 (M2): Density correction

An alternative approach is to adapt contact patterns to different demographic structures correcting by the density of available contactees, which we formalize with the following equation:

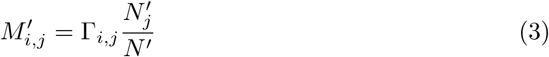

Thus, we interpret that the matrix *M_i,j_* is the product of two factors:

- The intrinsic connectivity matrix: Γ_*i,j*_
- The fraction of individuals in *j*: 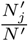

Thus, we are assuming that an individual has an intrinsic preference over certain age-groups depending on its age, captured by Γ_*i,j*_ and the final contact rate is modified according to the density of available contactees.

The matrix Γ_*i,j*_ corresponds, except for a global factor, to the contact pattern in a “rectangular” demography (a population structure where all age groups have the same density). We can obtain these matrices Γ*_i,j_*, that are country-specific, from survey data using equation 3:

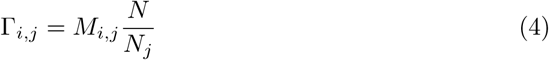

which allows to rewrite equation 3 as a function of the original matrix *M_i,j_*:

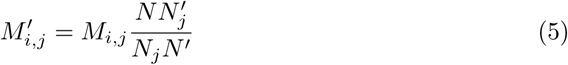

This methodology for adapting contact patterns has already been used by De Luca and collaborators, introducing the matrix Γ*_i_*,_*j*_ in the force of infection [8]. Also a similar correction is used in Prem et al. [28] to adapt European contact matrices to other countries (although this work integrates more data beyond demographic structures).

### Method 3 (M3): Density correction + Normalization

A cardinal feature of M2 is that it does not preserve the mean connectivity of the entire network of contacts. As a result, depending on the initial contact matrix and the dynamics of the demography, the evolution of the contact structure can produce average connectivities that depart strongly from its initial value. For the sake of disease modeling, this situation is essentially irrelevant if the contact rate of the outbreak to model can be callibrated at its early stages (i.e. its reproductive number). In that case, any global scaling factor multiplying the contact matrix is absorbed by the estimation of a larger or smaller infectiousness *β.* However, if that is not the case and epidemiological parameters measured in the past (i.e. a pathogen’s infectiousness) are used to generate forecasts of independent outbreaks that might occur later in time, the overall scaling factor of the contact networks become extremely relevant. In such scenario, to couple an a-priori characterization of a pathogen’s infectiousness on top of contact networks with different mean connectivities will artificially inflate or shrink the size of modelled epidemic events as a function of time. Although considering an evolution of the mean connectivity as demography changes might be reasonable, the inability of M2 of producing contact matrices of stable mean connectivities might suppose a liability in some scenarios.

Taking that potential issue into consideration, we have proposed an alternative approach that, in addition of correcting for the densities of contactees, preserves the mean connectivity of the overall system across time. Thus, an evolution of the mean connectivity could always be forced by adding a global factor in a controlled way.

To do so, we begin by defining 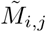 as the connectivity matrix from M2:

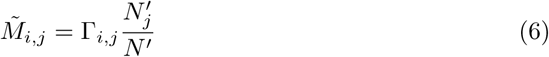

and then we divide it by its connectivity:

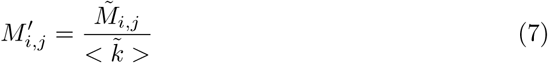

Thus:

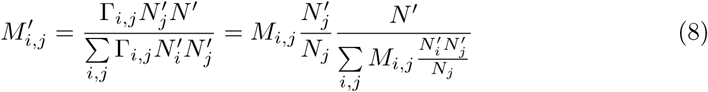

Notice that all methods trivially coincide in the year in which the data was obtained (i.e. when the survey was done). Also the definition of Γ*_i,j_* does not change between M2 and M3 in these cases, as the initial *M_i,j_* has been normalized to have a mean degree of 1, and we extract it with the same equation as before (eq. 4).

## Results

### Reciprocity error

In order to study the error incurred when using M0, we transform the contact matrices obtained from empirical studies in different countries to new matrices that correspond to the same location but at different years (that could be past records or future projections). As the population changes over time, the new matrices incorporate the population demographies of the same countries across time. We define the reciprocity error as the coefficient of variation of the number of contacts measured in both directions, which gives us a matrix that we will call non-reciprocity matrix (*NR_i,j_*). It is an antisymmetric matrix, in which a positive value of the entry (*i*, *j*) means that there are more contacts from *i* to *j* than in the opposite direction, and viceversa. A value of 0 would mean that the contacts between *i* and *j* are well balanced. More details can be found in the Supplementary Information.

In Figure 1 we represent the demographic structures of Poland (panel A) and Zimbabwe (panel B) for different years alongside the corresponding non-reciprocity matrices. In the case of European countries (Poland in panel A as an example), demographic structures have suffered from an ageing process during the last decades, which is predicted to continue in the future. This ageing tends to provoke negative values under the diagonal for the matrices *NR_i,j_* when we assumed past demographic structures, while the opposite will occur in the future. The behaviour for African countries (Zimbabwe in panel B) is slightly different, as their demographies have been more stable for the last decades and only now they are beginning to age faster. In brief, when we use directly a contact pattern in a demographic structure that is younger than when it was measured, it will lead to an overestimation of the contact rate of (and the force of infection corresponding to) the youngest age-groups. The opposite will occur when we use contact patterns in an older population.

**Fig 1.**
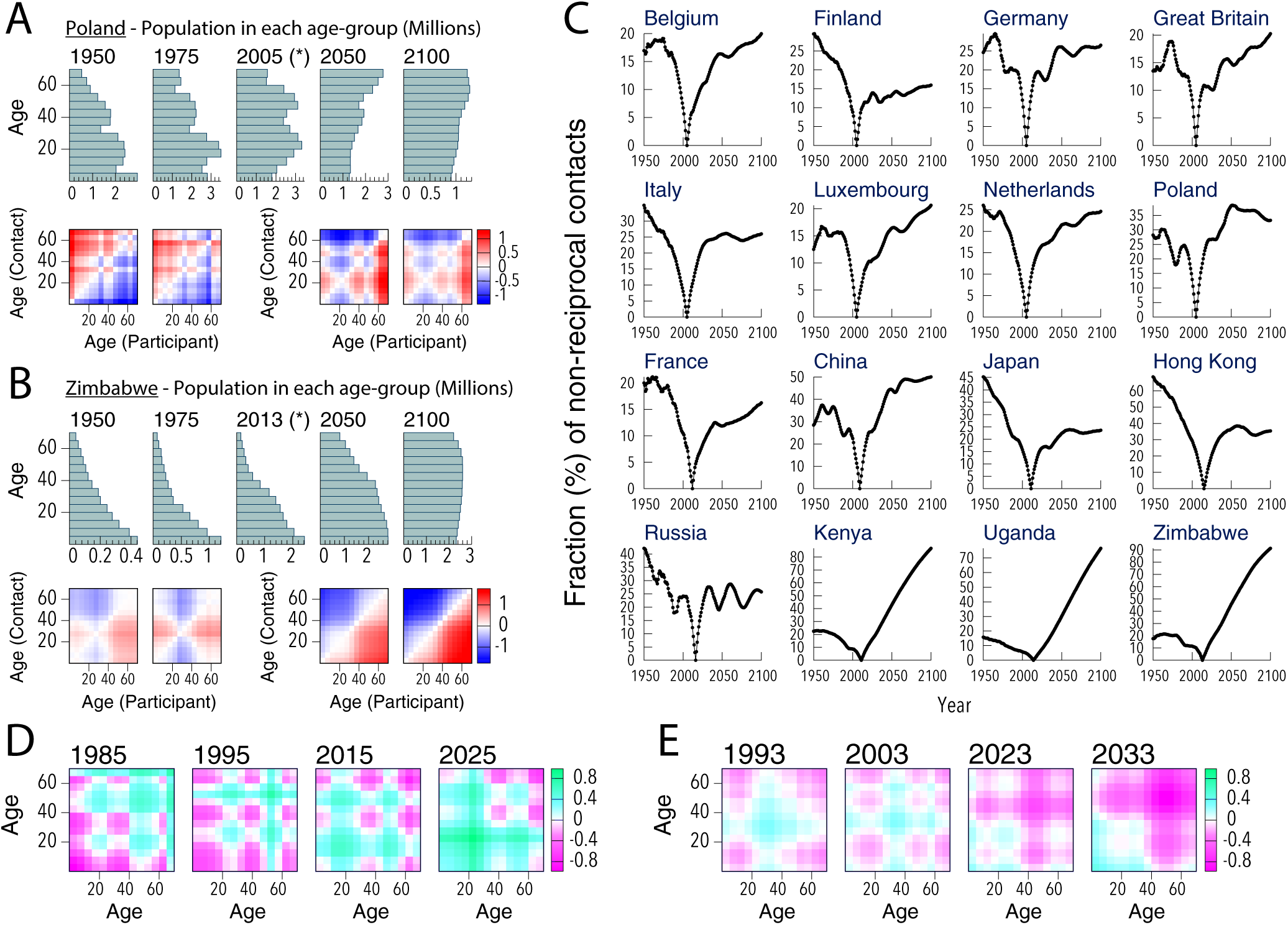
Analysis of methods M0 and M1. A-B: Demographic structures for different years and the respective non-reciprocal matrices 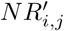 for Poland and Zimbabwe respective using M0. C: Evolution of the total fraction of non-reciprocal contacts for M0 in the 16 countries analyzed in this study. D-E: 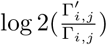 for Poland and Zimbabwe respectively, in four different years (10/20 years before/after the measurement of the contact patterns) for M1. The original data corresponds to 2005 for Poland and 2013 for Zimbabwe.

In figure 1C we represent the evolution of the proportion of non-reciprocal contacts for all 16 countries (see Supplementary Information). This magnitude is equal to zero in the year when the contact matrix was measured, as we have applied a correction for the empirical matrices to fulfill reciprocity at the reference setting. However, it dramatically increases as we move far from the year of the survey. In the examples shown here, only two years before/after the survey time, the fraction of non-reciprocal contacts already reaches 5%. Note that methods M1, M2 and M3 are well balanced by construction, thus *NR_i,j_* = 0 for every (*i, j*) when using any of them.

### Intrinsic Connectivity error

We next study the evolution of the ratio between the age-dependent contact rates and an homogeneous mixing scenario. This ratio gives us the matrix Γ*_i_*,_*j*_ defined as the intrinsic connectivity in equation 4. The entries of Γ*_i_*,_*jj*_ are bigger than 1 when the interactions between age-groups *i* and *j* surpasses what it is expected from the case of homogeneous mixing, and smaller than 1 in the opposite case. See the Supplementary Information for more details.

In Figure 1D and 1E we show 4 snapshots of the ratio of the intrinsic connectivity and the original survey 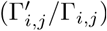 obtained using M1 for Poland and Zimbabwe respectively. Each panel corresponds to an adaptation of the contact matrix to the population demography of the countries 10 and 20 years before and after the survey (i.e., the 4 matrices correspond to *t* = *τ* − 20*y*, *t* = *τ* − 10*y*, *t* = *τ* + 10*y* and *t* = *τ* + 20*y*). We can see that, even if M1 corrects the appearance of non-reciprocity, this method changes the tendency of some age-groups to mix with respect to others. Specifically, we can see that M1 will over-represent contacts between young individuals (and under-represent contacts between old individuals) as the population gets older.

Furthermore, the previous results are quantitatively important. For instance, if we were to use the contact matrices that we have from Poland (measured in 2005) today (2018), we would have that the ratio 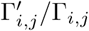 surpasses 1.5 for some specific age-group pairs, while it goes down to almost 0.5 in others, or, in other words, the usage of M1, which does not take into account the changes in the fractions of individuals in each age-strata that occurred between 2005 and 2018, causes a bias of more than 50% in the contact densities projected between certain age groups. Consequently we say that M1 does not preserve intrinsic connectivity. The density correction (M2) avoids this problem, as it explicitly considers a fixed intrinsic connectivity matrix (Γ*_i,j_* as defined in the Methods section) that is modified according to the density of each age-group (see equation 3).

### Evolution of mean connectivity

In Figure 2A-B we represent the contact patterns obtained with M2 and M3 for Poland and Zimbabwe, respectively, in different years. We see how, specially in the case for Zimbabwe, as the population gets older, the values of the matrix below the diagonal (contacts toward young individuals) fade in favor of contacts toward older individuals as those age-groups gain more representation. As for the mean connectivity (Figure 2C), considering the evolution of contact patterns in M2 or considering them constant (M0) leads to the same qualitatively behaviour, although variances are higher with M2. These trends are decreasing in Europe and increasing in Africa. M0 and M1 have the same mean connectivity, as M1 consists basically of a rewiring of those connections that exist in M0 in order to correct for reciprocity. M3 is a normalization of M2 so the connectivity is constant in this case.

**Fig 2.**
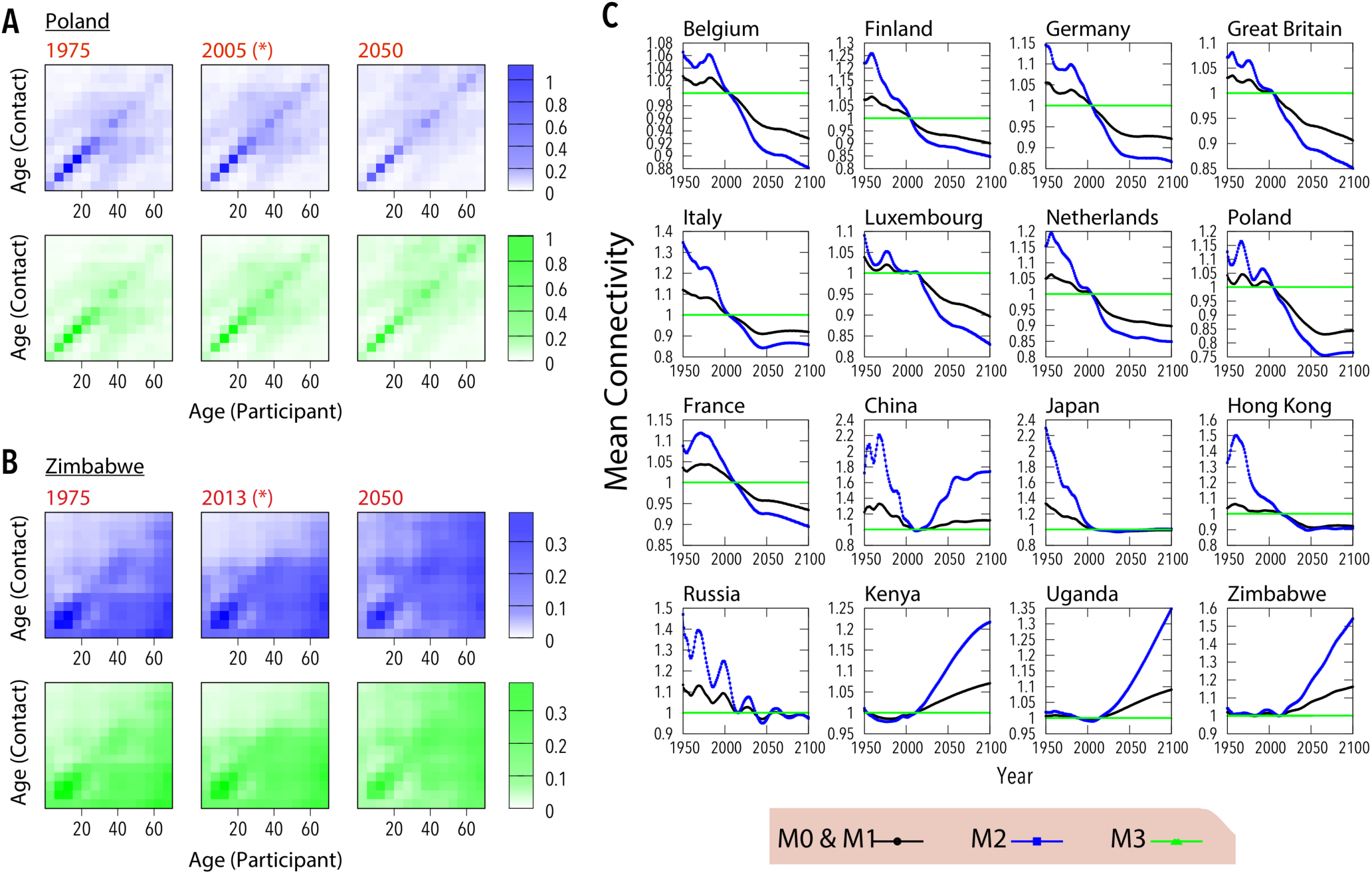
Analysis of methods M2 and M3. A-B: Contact patterns *M_i,j_* (*t*) for five different years with methods M2 (blue) and M3 (green) for Poland and Zimbabwe, respectively. C: Evolution of Mean Connectivity for M2 (blue), M3 (green) and M0 and M1 (black, both methods give the same mean connectivity).

### Overview of different methods

We have shown up to four different methods of use heterogeneous contact patterns when demography evolves in time (being the first one of them to simply use them without any further consideration regarding the demographic structure). In table 1 we summarize their properties.

**Table 1.** Properties of different methods

| Method | Reciprocity? | Preserves Intrinsic Connectivity? | Constant average connectivity? |
| --- | --- | --- | --- |
| M0: Unadapted contact patterns | No | No | No |
| M1: Pair-wise correction | Yes | No | No |
| M2: Density correction | Yes | Yes | No |
| M3: Density correction + Normalization | Yes | Yes (with a global factor) | Yes |

Summary of the different methods to deal with contact patterns and their properties.

### Geographical Comparisons

The intrinsic connectivity matrices Γ_*i,j*_ that we obtain for every country allow us to compare the contact patterns of different settings once the influence of demography has been accounted for, and removed. In Figure 3A we represent these matrices for the 16 countries analyzed in this work. Just by visual inspection we can identify some distinctive features: European matrices are more assortative and present higher interaction intensities among young individuals than African ones. To formalize this observation, in Figure 3B, we place the different matrices in a two dimensional plot defined by the proportion of overall connectivity produced by young individuals and the assortativity coefficient (see Supplementary Information for the definition of these quantities). African and European countries cluster around different values of these two magnitudes: specifically, in African countries we found less assortativity and the contacts are less dominated by young individuals than in the European countries. As for the Asia region we see that Japan and China have significantly higher assortativity and fraction of contacts among young individuals than either African or European countries. In turn, Hong Kong, with its particular geographic idiosyncrasy- a small country, predominantly urban, with one of the highest population densities in the world-, presents an intrinsic connectivity matrix that is more similar to one from a European country than from China or Japan.

**Fig 3.**
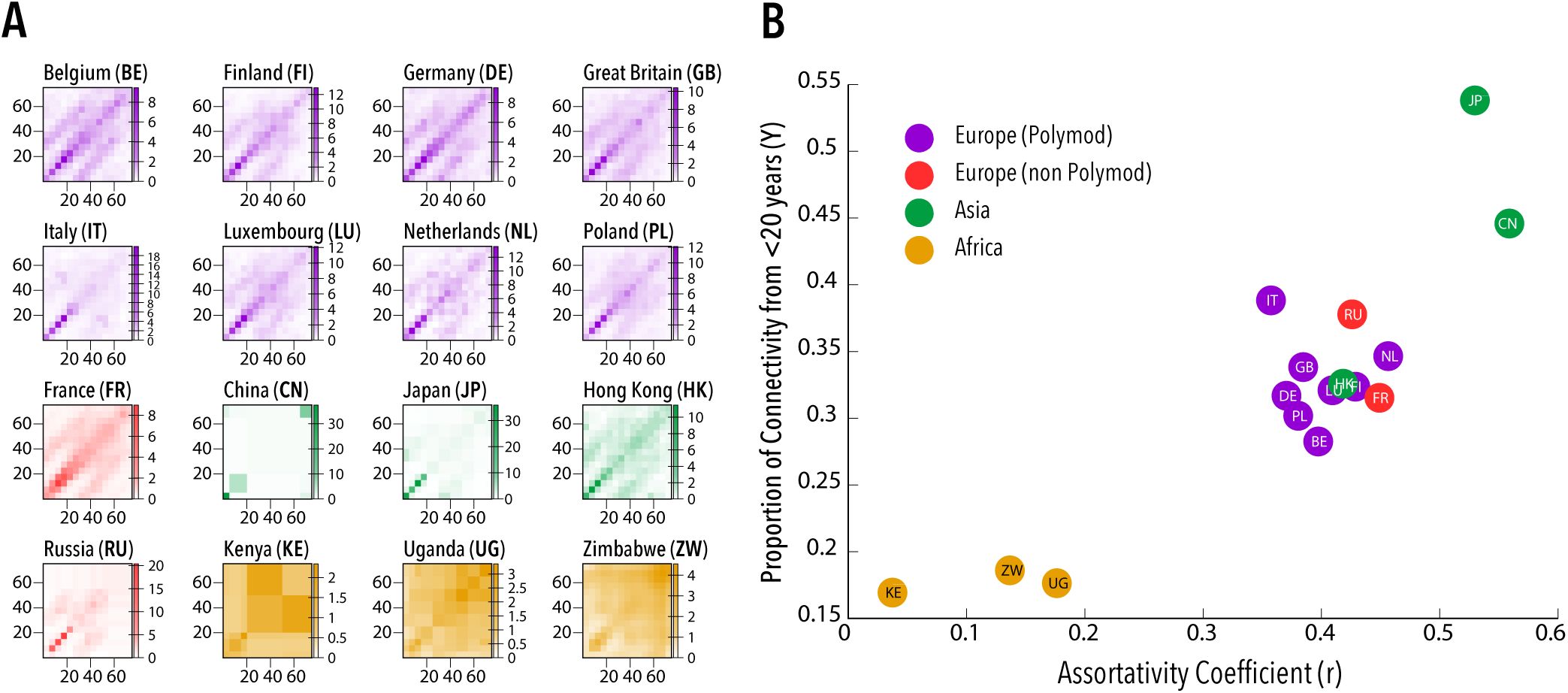
Geographical comparison of empirical contact matrices. A: Γ*_i,j_* matrices for the 16 countries considered in this work. B: Proportion of the overall connectivity that comes from individual with less than 20 years (Y) vs the assortativity coefficient (r) for the 16 countries.

### Short cycle SEIR dynamics

Up to now, we have shown that there are several ways to deal with demographic change and evolving populations regarding the structure of the contact patterns for a given population. We next address how these different methods impact disease modeling. To this end, we implement a Short cycle SEIR model (details can be found in the Supplementary Information) to study a situation where a short-cycle, influenza-like pathogen appears in a given location, at different possible times, associated to the same reproductive numbers. Under this hypothetical scenario, we would like to know how different would be the forecasted size of the epidemic as a result of considering different contact matrices coming from the different projection methods proposed in this work. In particular, this scenario is instrumental to distinguish the outcomes from models M0,M1 and M2. However, the requirement of the outbreaks to have the same reproductive numbers implies the assumption that the infectiousness *β* can be estimated independently in each event. As a consequence, since the matrices derived from M2 and M3 only differ by a global scaling factor, this operation absorb the differences between M2 and M3, making them indistinguisable.

The results of this exercise are presented in Figure 4. In Figure 4A we can see that, while methods M0 and M1 predict lower age-aggregated incidences in European countries in 2050 with respect to 2000, M2 reduces these differences and the incidences are comparable for both years or even positive. A different situation occurs in Africa, where M0 and M1 predict an increase in incidence in the future while using M2 would lead to a decrease, though differences remain small (less than 5% of variation).

**Fig 4.**
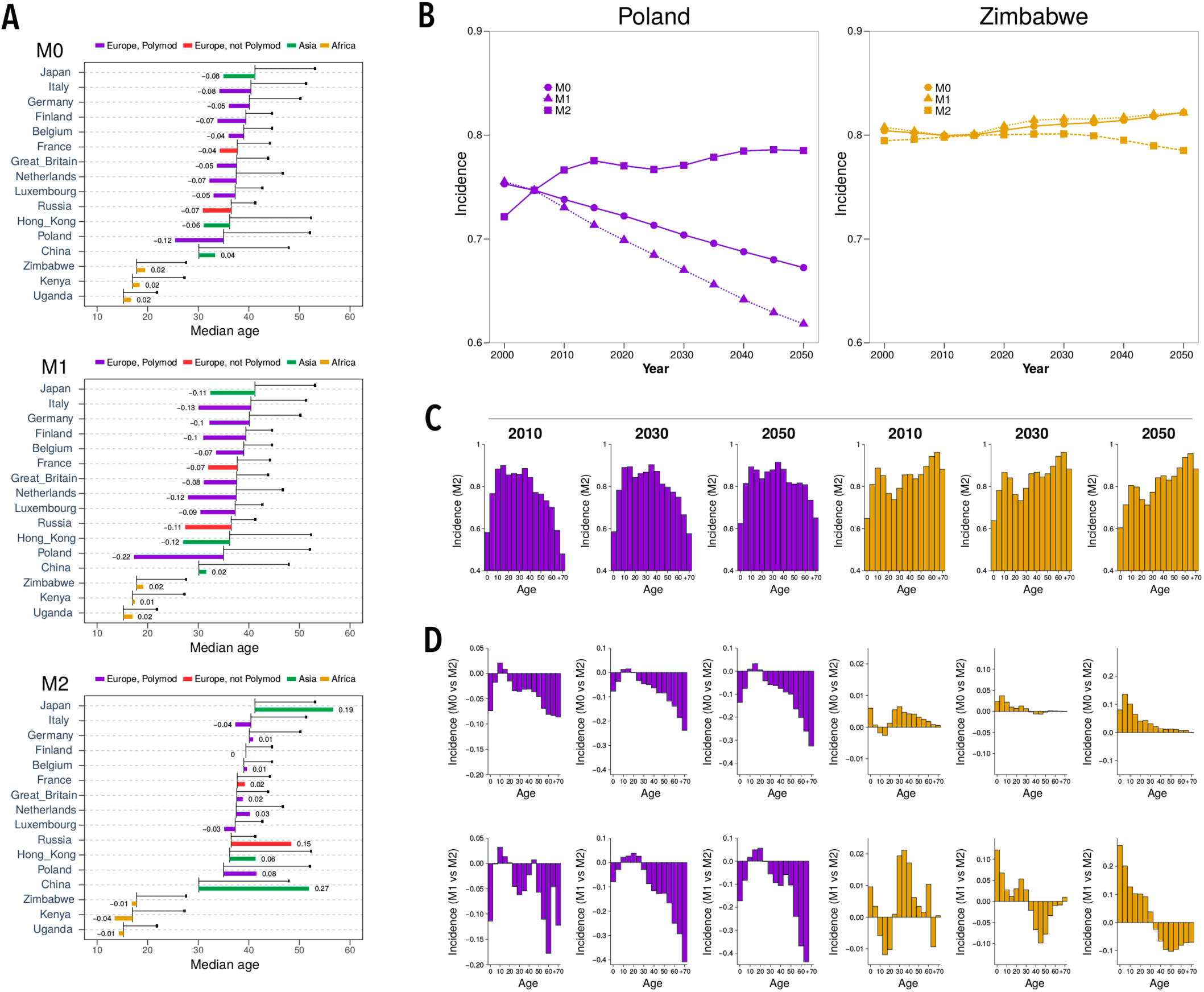
SEIR dynamics. A: Median age at 2000 and 2050 (black line, beginning with the value at 2000 and ending with a bullet point with the value at 2050) for the 16 countries considered and relative variation in incidence over the same period (colored bars), for M0, M1 and M2. B: Incidence (over all ages) vs Year for Poland (blue) and Zimbabwe (orange) using M0, M1 and M2. C: Incidence by age group for Poland and Zimbabwe in 2010, 2030 and 2050 using M2. D: Relative differences of the incidence by age group of M0 and M1 with respect to Me (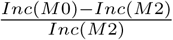 and 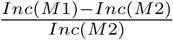).

In panel 4B we represent, for two examples of Europe and Africa (Poland in blue and Zimbabwe in orange), the temporal evolution of the incidence observed with the different methods. Furthermore, we represent the age-specific incidence for both countries in three different years: 2010, 2030 and 2050 (Panel 4C). The age-distribution of the incidence evidences the differences in connectivity patterns between Poland and Zimbabwe. While the incidence in elderly people drops in Poland (as the contact rates for those age-groups also drop), it remains high in Zimbabwe for the same age-groups.

The different methods of implementing contact rates also affect the age-specific incidence. In panel 4D we represent the relative variation in age-specific incidence obtained with methods M0 and M1 with respect to M2 for Poland and Zimbabwe. In Poland we see that M0 and M1 tend to underestimate the incidence specially among the elder age-groups. In Zimbabwe M0 tends to overestimate the incidence among young individuals, while with M1 we encounter both effects: and overestimation among the youngest and a underrepresentation among the eldest.

The reshaping of the age-specific incidence between models is coherent with the changes in topology already studied. For the case of M0, i.e., maintaining the contact patterns constant in time, we have that in the future, as the demographic structure shifts to older populations, contacts toward children will be overrepresented and contacts toward adults will be underrepresented. At first order we can obviate the contacts that are far from the diagonal, and establish that M0 mainly underrepresents contacts between adults and overrepresents contacts between young individuals (in the context of aging populations). Thus, we will obtain an underrepresentation of the incidence in adults, and the opposite in children. However, as the eldest age-groups increase their population in Europe, they dominate the dynamics and cause and underestimation of the global incidence that eventually affects all age-groups. In African countries, where the contact patterns are less assortative than European countries, this effect is smaller. Besides, as African populations are still young even in 2050, the overestimation of young contacts dominates the dynamics, and the differences in incidence are mainly positive. The situation is similar for M1. As represented in Figure 1D-E, for M1 we also have an underrepresentation of contacts between adults and an overestimation between young individuals, yielding to similar results than M0.

All together, these results illustrate how an ill adaptation of the contact patterns observed in the past in a given country to a later time point can translate into epidemiological forecasts that are highly biased. Regarding the dynamic equivalence of methods M2 and M3, we have to emphasize that it emanates only from the assumption that reproductive numbers can be measured at the early stages of any of the epidemics being simulated in each year, which is a conservative -often optimistic- assumption. Alternatively, we could think of an scenario where the reproductive number of a given pathogen was estimated in a given year, and that information used to infer the probability of transmission per contact (the infectiousness *β*) of the pathogen, with the aim of producing a-priori forecasts for posterior re-apparences of the same pathogen. In such alternative scenario, the usage of different contact matrices projections would be even more relevant, for it would impact directly the reproductive number of the forecasted outbreaks, now characterized by a common *β.* In such an scenario, (which is conceptually similar to the task of producing long term forecasts of persistent diseases [12], based on epidemiological parameters calibrated on an initial time-window), the dynamic equivalence of models M2 and M3 is broken, since running M2 or M3 with the same infectiousness parameter and different average connectivities yields different reproductive numbers.

## Discussion

Summarizing, empirical contact patterns belong to a specific time and place. If we want to integrate the heterogeneity of social mixing into more realistic models, we need to address how (and in what cases) to export contact patterns from empirical studies to the populations we want to study. In this work, we have studied and quantified the significant bias incurred when a specific contact pattern is blindly extrapolated to the future (or the past), even if we remained inside the same country where those contacts were measured. In fact, only a couple of years after the measurement of these contact patterns, the changes in the age structure of the population make them vary significantly. Thus, for any meaningful epidemic forecast based on a model containing age-mixing contact matrices, we would need to adapt them taking into account the evolution of the demographic structures. Moreover, as we have shown, even for cases that do not expand into long periods of time and a constant demography could be assumed, it is necessary to make an initial adaptation of whatever empirical contact structure we want to implement, into the specific demographic structure of our system. We have also seen how these relevant differences in the topology of contacts yield to significant consequences for the spreading of a disease. Applying different methods to deal with contact patterns leads to important differences not only in the global incidence for a SEIR model, but also on age-specific incidences. Having such an important impact for the spreading of a disease, the insights provided by this work should be taken into consideration by modelers and also by public health decision-makers.

In a similar way, we have explored the differences between the contact patterns of different countries. Thus, we have found the existence of some specific characteristics beyond the underlying demographic pyramid, which warns against exporting contact patterns across different geographic areas (i.e. continents). As there exists different intrinsic connectivity patterns (i.e., once demography effects have been subtracted) between countries, it is also likely that there exists a time-evolution of the intrinsic connectivity inside the same setting. Although it is impossible to predict how society will change in the future, we should always take this into account as a limitation in any forecast for which the heterogeneity in social mixing is a key element.

Finally, we note that there are some limitations that could affect quantitatively the results shown in this work. First of all, we have derived the contact patterns of the different studies according to the demographic structures of the specific country for the year the survey took place. Thus, we are implicitly assuming that the setting where the different surveys were performed are comparable with the national data in terms of their demographic pyramids. In other words, we assume that the surveys are representative of the population at large. This is likely true for most of the countries analyzed, but there are certin cases in which this might not be the case, either because of small study size or putatively biased recruitment of participants. Besides, as we have already discussed in the Methods section, the different granularity (i.e., definition of the age-groups) used throughout the bibliography studied also imposes some limitations when comparing the data. It is also worth pointing out that, although in this work we have focused on age-structured systems (which has had its relevance in recent history of epidemiology), the problem studied here can be extrapolated to other models that might categorize their individuals based on other different traits that determine their social behavior.

The results reported here and their implications open several paths for future research. One is related to the social mixing patterns themselves. In order to predict the large-scale spreading of a disease, multiple scales need to be integrated and coupled together. This implies that when integrating different spatial scales, we need to deal with different contact matrices and local demographies. For instance, in developed countries, it is known that the structure of the population is not the same in the most central or most populated cities as compared to smaller ones or the countryside. Thus, nation-wide demographies and surveys to infer contact matrices might need to be disaggregated. What is the right spatial scale to measure both quantities is an interesting and unsolved question. In this sense, here we have limited our simulated disease scenario to the case of isolated populations –a country–, but it remains to be seen what are the effects over a meta-population framework, in which we have mobility between subpopulations of potentially very different demographic structures. We plan to explore these issues in the future.

## Supporting information

**S1 Supporting Information** Extended details on methods and additional analyses.

## Acknowledgments

We thank V. Colizza for helpful comments regarding the implementation of the SEIR model. S.A. is supported by the FPI program of the Government of Aragón. A.A is supported by an FPI doctoral fellowship. J.S. is supported by the Canadian Institutes of Health Research through a Banting fellowship. This work was partially supported by by a grant to the group FENOL, by MINECO through grant FIS2017-87519-P.

